# Diversity of *Escherichia coli* from the enteric microbiota in patients with *Escherichia coli* bacteraemia

**DOI:** 10.1101/561613

**Authors:** Mia Mosavie, Oliver Blandy, Elita Jauneikaite, Isabel Caldas, Matthew J. Ellington, Neil Woodford, Shiranee Sriskandan

**Author notes:** Correspondence: Shiranee Sriskandan, Imperial College London, Hammersmith Campus, Du Cane Road, London W12 0NN, UK.

## Abstract

**Objective:** The increase in *Escherichia coli* bloodstream infections mandates better characterisation of the relationship between commensal and invasive isolates. This study established a simple approach to characterize diversity of *E. coli* in the gut reservoir from patients with either *E. coli* bacteraemia, other Gram-negative bacteraemia, or patients without bacteraemia not receiving antibiotics. Stool or rectal swabs from patients in the three groups were cultured on selective chromogenic agar. Genetic diversity of *E. coli* in gut microbiota was estimated by RAPD-PCR.

**Results:** Enteric samples from *E. coli* bacteraemia patients yielded a median of one *E. coli* RAPD pattern (range 1-4) compared with two (range 1-5) from groups without *E. coli* bacteraemia. Of relevance to large-scale clinical studies, observed diversity of *E. coli* among *E. coli* bacteraemia patients was not significantly altered by sample type (rectal swab or stool), nor by increasing the colonies tested beyond ten. Overall, hospitalised patients demonstrated an apparently limited diversity of *E. coli* in the enteric microbiota and this was further reduced in those with *E. coli* bacteraemia. The reduced diversity of *E. coli* within the gut during *E. coli* bacteraemia raises the possibility that dominant strains may outcompete other lineages in patients with bloodstream infection.

## INTRODUCTION

The increasing burden of *Escherichia coli* bacteraemia in the UK and dominance of antimicrobial-resistant lineages worldwide [1–4] point to a need for better understanding of the reservoir and diversity of *E. coli* in the gut microbiota of those affected by invasive *E. coli* infections. *E. coli* is present in abundance in the adult gut microbiota, however the diversity of *E. coli* strains within the gut reservoir has only been studied in limited groups that do not include bacteraemia patients [5–10].

The enteric microbiota of such patients are hard to study, as bacteraemic patients will be acutely unwell, and stool samples are usually not readily available or clinically indicated, albeit that routinely-collected samples could represent an accessible sample source. Patients with suspected bacteraemia will usually receive empirical antimicrobial therapy within an hour of admission to hospital, consistent with international guidance on sepsis [11], thus all enteric samples are potentially antibiotic-exposed and may also be refrigerated for long periods of time before reaching the diagnostic laboratory, in turn potentially affecting microbiota [12].

Randomly Amplified Polymorphic DNA (RAPD) analysis can provide a relatively crude molecular method to differentiate strains based on amplicon banding patterns; while such data are not directly comparable between studies, the method can be used to assess genetic diversity of *E. coli* isolates rapidly and cost-effectively [8, 13, 14]. The aims of this work were to establish simple methods to characterize the diversity of *E. coli* strains present within the intestinal microbiota of hospital patients with *E. coli* bacteraemia; to evaluate control hospitalised groups; and to assess scalability and establish a sample collection for future large-scale genomic studies.

## METHODS

### Patients and Samples

Enteric samples were stool or rectal swabs from hospital in-patients obtained within 72h of onset of Gram negative bacteraemia in west London NHS Teaching Hospital sites between 1^st^ July 2015 and 4^th^ August 2016. Enteric samples were sought from patients prior to final microbiological identification; groups were subsequently categorised into confirmed *E. coli* bacteraemia or other Gram negative non-*E. coli* bacteraemia. Controls were selected from inpatients within the same hospital wards that had not received an antibiotic in the preceding 30 days. Samples were obtained with informed consent except where rectal swabs had already been submitted to the hospital diagnostic laboratory for routine screening. In total, the study included 462 enteric samples (Figure S1). Samples were refrigerated and analysed within 48h unless otherwise stated, after any routine diagnostic testing had been completed. Preliminary studies demonstrated that refrigeration in the research lab for 48-72h had little, or no, effect on the retrieval of *E. coli* from enteric samples though yield reduced by 20-25% over 10 days (data not shown). The study was approved by a research ethics committee (Reference 14/LO/2217).

### Culture methods

Approximately 10-60 mg of stool was suspended in 3 ml 0.9 % saline to achieve an optical density of A_600_ 0.1 [15]. The suspension was diluted 100-fold in 0.9% saline and 100 µl (~300-500 CFU) cultured on coliform-selective chromogenic agar (Brilliance™ *E. coli* coliform Selective Agar, Oxoid, UK) and Columbia Blood Agar (CBA) (EO Labs, UK) at 37°C overnight in 5% CO_2_. Samples with low counts (0-5 CFU) after 24h growth were re-plated using 100 µl of undiluted suspension. Rectal swabs were plated directly on agar as described above. To confirm presence of *E. coli* in a selection of stool samples, DNA was extracted directly from stool using Qiagen QIAamp DNA Stool Extraction Kit (Qiagen, UK) and *gadA* PCR was performed as previously reported [16] (Table S1). Samples were separately plated and all colonies frozen in glycerol bead stocks for future use.

### Evaluation of colony number required to assess diversity

To determine whether the number of *E. coli* colonies tested per plate would significantly impact on the measured diversity of enteric *E. coli*, groups of 5, 10, 15 or 20 *E. coli* single colonies were selected from individual samples that had been cultured on Brilliance agar. Samples were from randomly selected hospitalised patients with *E. coli* bacteraemia receiving antibiotics (n=5), and from randomly selected hospitalised patients who did not have bacteraemia and were not receiving antibiotics at the time (n=5). Individual colonies were boiled at 90 °C for 10 min to lyse the cells and release DNA; these were then subject to Randomly-Amplified-Polymorphic-DNA (RAPD)-PCR using primers and methods previously reported [17] (Table S1). The number of different RAPD patterns obtained from 5, 10, 15 or 20 single *E. coli* colonies was compared for each sample.

### Comparison of *E. coli* diversity between patient groups

Single purple colonies from chromogenic agar identified as *E. coli* were randomly selected from different quadrants of the chromogenic agar plate, and sub-cultured on to LB agar (Oxoid, UK). Ten colonies per sample were subject to DNA extraction and RAPD-PCR as described above. The number of different RAPD patterns obtained was compared for each of the patient groups. Groups were compared pairwise by Mann-Whitney test; groups of more than two were compared by Kruskall-Wallis (Graph Pad Prism)

## RESULTS and DISCUSSION

### Number of colonies necessary to assess diversity of *E.coli* isolates in enteric microbiota

Although up to 10 distinct *E. coli* lineages can be present in the normal human enteric microbiota, medians of 2-3 types have been reported in patients in the community [6–10]. For a large clinical study in bacteraemia patients, that might include 100-200 hospitalised cases, exhaustive molecular analysis of 1000’s of *E. coli* colonies would not be feasible. To determine a suitable number of *E. coli* colonies to analyse per patient when evaluating the diversity of *E. coli* lineages in the gut microbiota of hospitalized patients, we compared RAPD patterns of 20, 15, 10 and 5 *E. coli* colonies cultured from ten different in-patient faecal samples. Five patients had been diagnosed with *E. coli* bacteraemia and were receiving antimicrobial treatment, while five patients were non-bacteraemic in-patients and had not been receiving any antibiotics for 30 days prior to sample collection.

Overall, a median of two RAPD patterns was identified per sample; however *E. coli* bacteraemia patients had a reduced range of RAPD types (median one, range 1-3) compared with non-antibiotic exposed patients (median two, range 1-4). Examination of 15 colonies allowed detection of increased diversity in just one sample tested in each group compared with examination of 10 colonies. Further expansion to examine 20 colonies did not increase the observed diversity of *E. coli* per sample in either group. Overall there was no difference in the number of potential different genotypes detectable by RAPD-PCR when selecting 10 or 20 colonies (P >0.05) (Figure 1 and Table S2). As such, for subsequent work, a practical approach of selecting 10 colonies per patient was adopted.

**Figure 1.**
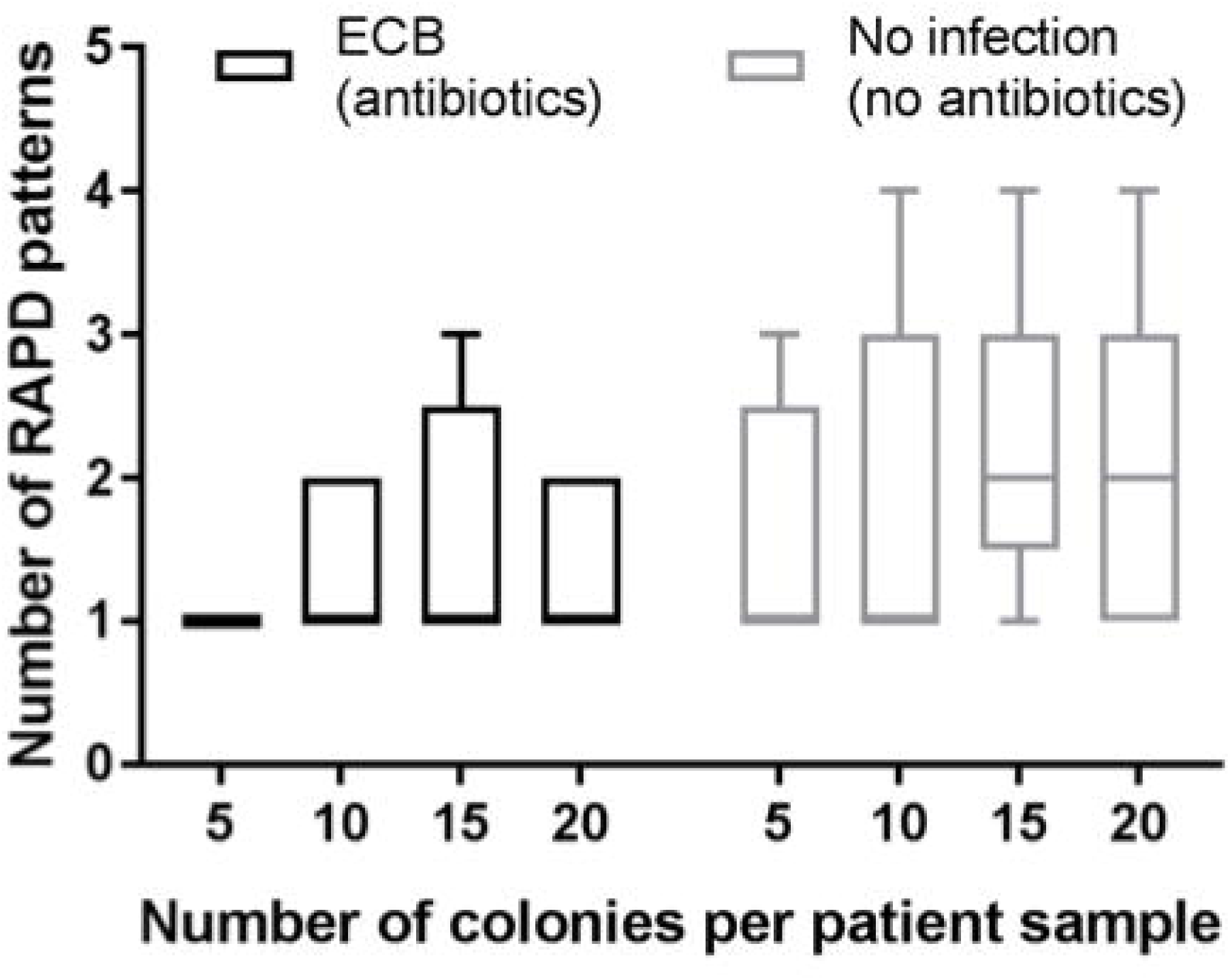
Number of RAPD patterns detected when testing 5, 10, 15 or 20 *E. coli* colonies per patient sample. Enteric samples were tested from *E. coli* bacteraemia (ECB) patients (n=5, receiving antibiotics), and non-antibiotic-exposed inpatient controls (n=5, no infection, not receiving antibiotics). Different numbers of colonies were evaluated, as indicated and the number of distinct RAPD patterns enumerated. Plot shows median and interquartile range.

### Assessment of *E. coli* diversity between patient groups

*E. coli* diversity was then compared between larger sets of enteric samples from *E. coli* bacteraemia patients (receiving empiric antibiotics) and control in-patients not receiving antibiotics. To control for the potential effect of antibiotic exposure we included an additional group of other Gram-negative (non *E. coli*) bacteraemia patients, who also received empiric antibiotics. Samples were sought from all eligible patients during the study period and RAPD-PCR was undertaken in all cases where enteric samples yielded at least 10 colonies of viable *E. coli* on subculture. (Figure S1)

The median number of *E. coli* RAPD patterns recovered from 70 hospital in-patient controls, who were not receiving antibiotics, was two (range 1-5), in contrast to the 74 *E. coli* bacteraemia patients in whom a median of one RAPD type (range 1-4) was detected (p=0.029). Despite being similarly antibiotic-treated, the median number of RAPD patterns detected in the 42 non-*E. coli* Gram negative bacteraemia samples analysed was two (range 1-5) although the differences between the three groups overall was not significant (p=0.06) (Figure 2).

**Figure 2.**
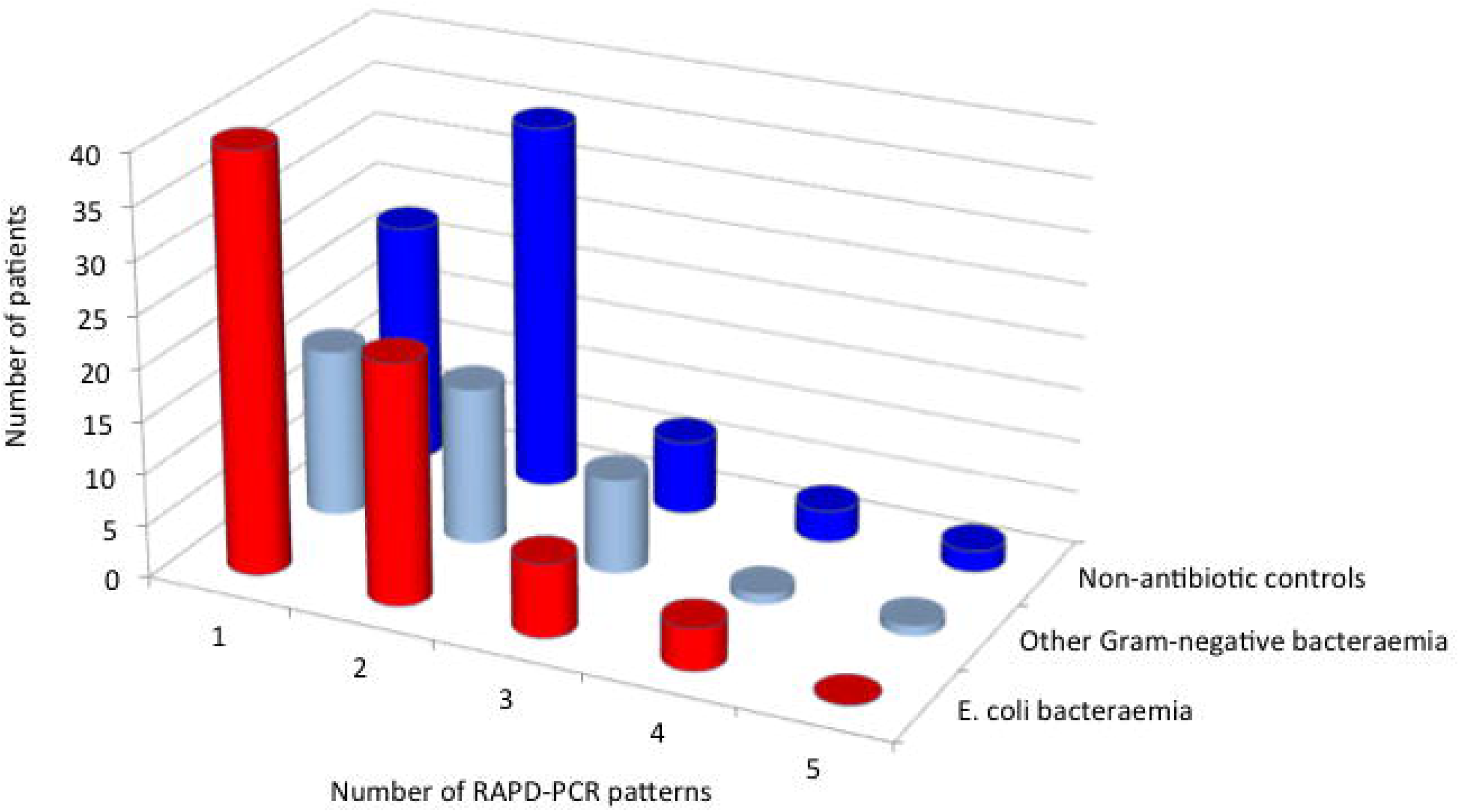
Diversity of *E. coli* in enteric microbiota from patients with *E. coli* bacteraemia compared with other patient groups. Frequency of different numbers of RAPD patterns among *E. coli* bacteraemia patients (n=74), patients with other Gram negative bacteraemia (n=42), and non-antibiotic exposed inpatient controls who had no infection and were not taking antibiotics (n=70). Although a difference was detected between the two main groups (*E. coli* bacteraemia and non-antibiotic inpatient controls, p=0.029), there were no overall differences between the three groups (p=0.06).

We considered whether the type of sample affected our results and so compared data obtained from rectal swabs and faecal samples; rectal swabs were performed largely for the purpose of screening for carriage of carbapenem-resistant organisms (CRO) using risk-based algorithms and could represent a marker of prior healthcare exposure. Importantly, however, there was no significant difference between the two types of sample (Figure S2). Rectal swabs may be obtained earlier during each hospital admission than stool samples, reducing the effect of recent antibiotic exposure on the diversity detected. The value of using rectal swabs is a useful observation, since inpatients are often unable to provide a faecal sample, while rectal swabs are increasingly obtained as part of routine screening practice.

Overall, we found a narrower diversity of *E. coli* in *E. coli* bacteraemia patients compared to control hospitalised patients. Although our study was not sufficiently powered to directly compare the *E. coli* bacteraemia group with the ‘other Gram negative bacteraemia’ group, our data suggest that the reduced diversity of *E. coli* in the enteric microbiota of *E. coli* bacteraemia patients may be specific to this group, and not necessarily related to concurrent antimicrobial exposure. We speculate that pathogenic *E. coli* strains causing bacteraemia may outcompete and dominate the gut microbiota. Whether this is related to past antimicrobial consumption or other pressures affecting the enteric reservoir of in-patients will require further study using specifically selected populations of patients with known antimicrobial history.

## LIMITATIONS

Our study had limitations, in that enteric samples were only available from around a quarter of eligible patients with bacteraemia, and, for practical reasons, efforts to culture *E. coli* from all patients did not include heat shock or novobiocin-enrichment which, in separate studies, we found did improve yield (not shown). However, the number of samples evaluated was high, and there is no evidence to suggest these limitations would have affected any one patient group more than another. The number of different *E. coli* lineages detected in the enteric microbiota of our control groups is similar to that reported by other investigators [8] although, to our knowledge, acutely unwell hospital inpatients have not been studied previously. RAPD-PCR can reliably investigate the diversity of *E. coli* subclones in enteric microbiota [14] although cannot be used to finely discriminate between isolates at the genomic level. It does however provide an affordable and scaleable method to determine the number of isolates to study using more expensive methods such as genome sequencing.

## Declarations

### Ethics approval and consent to participate

The collection and use of samples and consent process was approved by a research ethics committee (Reference 14/LO/2217).

### Consent for publication

Not applicable

### Availability of data and material

All data generated or analysed during this study are included in this published article [and its supplementary information files].

### Author contributions

MM, OB, and IC were responsible for patients and sample recruitment; MM, ME and EJ were responsible for sample analysis; MM and EJ were responsible for data analysis; NW and SS were responsible for project design and supervision; MM and SS wrote the first draft of the paper; all authors read and approved the final draft.

#### Acknowledgements

The authors thank the Imperial College NHS Trust Diagnostic Laboratory (Dr. Hugo Donaldson and colleagues and the BioAID team) for support and assistance with sample collection, and Dr. Kate Honeyford for advice on data analysis. SS acknowledges support of the NIHR Biomedical Research Centre awarded to Imperial College NHS Trust.

