## Supplementary Data for "Diversity of *Escherichia coli* from the enteric microbiota in patients with *Escherichia coli* bacteraemia"

**Supplementary Data File**

Table S1	Page 1

Table S2	Page 2

Figure S1	Page 3

Figure S2	Page 4

**Table S1 Primers used in this study**

| Primer |  | Sequence | Reference |
| --- | --- | --- | --- |
| **Detection of E. coli** | |  |  |
| *gadA* | Forward | 5' GATGAAATGGCGTTGGCGCAAG 3' | Doumith 2010 |
|  | Reverse | 5' GGCGGAAGTCCCAGACGATATCC 3' |  |
| **RAPD PCR** | |  |  |
|  | Primer 1 | 5’ GGTGCGGGAA | Radu 1998 |
|  | Primer 2 | 5’ AACGCGCAAC |  |

**Table S2 – Frequency of different RAPD patterns in each patient sample.**

|  |  | Number of colonies tested from each sample | | | |  |
| --- | --- | --- | --- | --- | --- | --- |
|  |  | 5 | 10 | 15 | 20 |  |
|  | Individual patient ID | Number of different RAPD patterns observed among *E. coli* colonies selected | | | | Maximum no. of RAPD patterns observed per patient sample |
| *E. coli* bacteraemia (receiving antibiotics) | A | 1 | 1 | 1 | 1 | 1 |
|  | B | 1 | 2 | 3 | 2 | 3 |
|  | C | 1 | 1 | 1 | 1 | 1 |
|  | D | 1 | 1 | 1 | 1 | 1 |
|  | E | na | 2 | na | 2 | 2 |
| Control inpatients  (not receiving antibiotics) | F | 1 | 1 | 1 | 1 | 1 |
|  | G | 1 | 1 | 2 | 2 | 2 |
|  | H | 1 | 1 | 2 | 1 | 2 |
|  | I | 2 | 2 | 2 | 2 | 2 |
|  | J | 3 | 4 | 4 | 4 | 4 |

na, not available as insufficient colonies from sample

**Figure S1	Sample recruitment for study.** Enteric samples were sought from every patient with confirmed Gram negative bacteraemia, provided the patient had been admitted to one of the participating hospitals. Enteric samples were only accepted if provided within 72h of bacteraemia onset. Samples comprised rectal swabs that were obtained as part of routine screening for carbapenem-resistant organisms or stool. Samples were initially screened for the presence of viable *E. coli* by plating on chromogenic agar without enrichment; purple colonies (10 per sample) were then subcultured onto LB agar and viable colonies tested by RAPD-PCR. The number of samples tested by RAPD-PCR was 74 (from 168 *E. coli* bacteraemia patient samples); 42 (from 144 non-*E. coli* Gram negative bacteraemia patients); and 70 (from 150 control in-patients not receiving antibiotics).

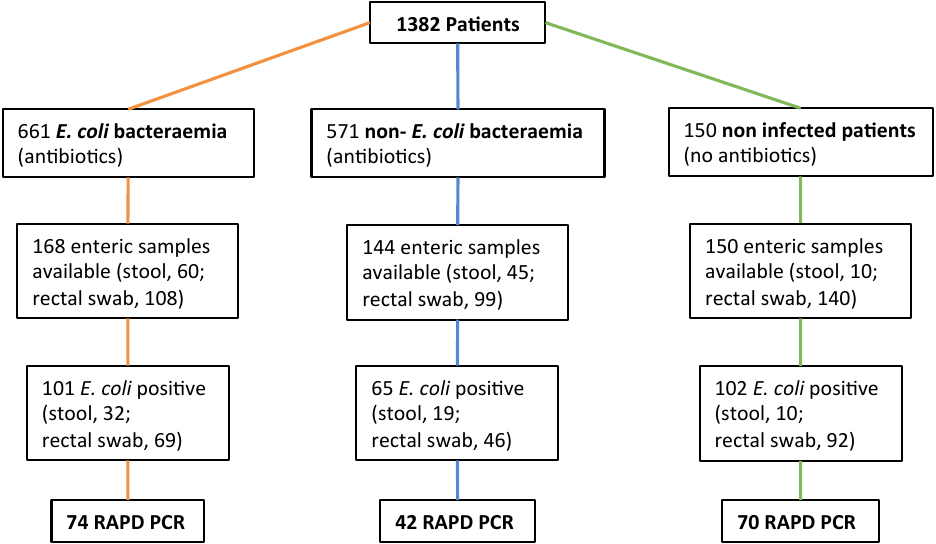

**Figure S2**	***E. coli* diversity in rectal swab and stool samples**. Number of RAPD patterns detected in each enteric sample from patients with *E. coli* bacteraemia (ECB); patients with other non-*E coli* Gram negative bacteraemias (non-ECB); and control in-patients (CON) who were not receiving antibiotics. Results for stool samples and rectal swabs are shown separately in each group. Plots show median (horizontal bar) and interquartile range. Differences between rectal swab and stool samples were not significant

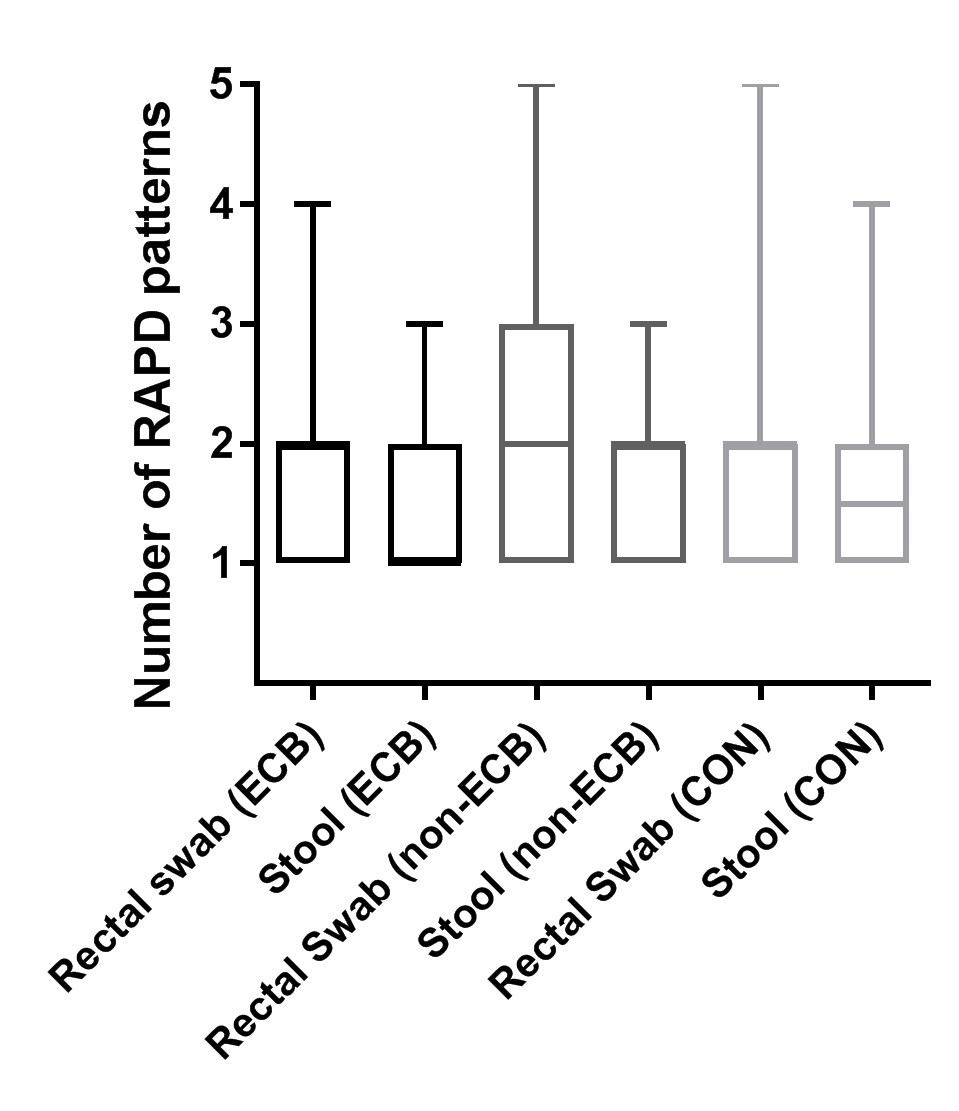
